# Eosinophils restrict diesel exhaust particles induced cell proliferation of lung epithelial A549 cells, vial interleukin-13 mediated mechanisms: implications for tissue remodelling and fibrosis

**DOI:** 10.1101/2021.03.30.436369

**Authors:** Rituraj Niranjan, Muthukumaravel Subramanian, D. Panneer, Sanjay Kumar Ojha

## Abstract

Diesel exhaust particulates (DEPs) affect lung physiology and cause serious damage to the lungs. A number of studies have demonstrated that eosinophils play a very important role in the development of lung tissue remodelling and fibrosis. However, the exact mechanism of its pathogenesis is not known. We for the first time demonstrate that, Interleukin-13 plays a very important role in the development of tissue remodelling and fibrosis. We demonstrate that, Diesel exhaust particle significantly induce eosinophils cell proliferation and interleukin-13 release in invitro culture conditions. Supernatant collected from DEP induced eosinophils cells significantly restrict cell proliferation of epithelial cells due to exposure of diesel exhast particles. Furthermore, purified interleukin-13 decreases the proliferation of A549 cells. Notably, Etoricoxib (selective COX-2 inhibitor) did not inhibit DEP-triggered release of interleukin-13, suggesting another cell signalling pathway. In, vivo exposer of DEP to the mice lung, resulted in the high level of eosinophils degranulation as depicted by the EPX-1 immunostaining and altered level of mRNA expressions of inflammatory genes. We also found that, a-SMA, fibroblast specific protein (FSP-1) has been changed in response to DEP in the mice lungs along with the mediators of inflammation. Altogether, we elucidated the mechanistic role of eosinophils in the DEP triggered proliferation of lungs cells thus providing an inside in the pathophysiology of tissue remodelling and fibrosis of lungs.

## 1. Introduction

Diesel exhaust particles (a component of environmental soot) are considered to produce many diseases in humans [1-3]. Diesel exhaust particles possesses chemical and physical properties which causes the toxicity in the body [3, 4]. Diesel enginines mainly vehicles are the main source of DEPs which contribute to the major percentage of particulate matter (PM), up to 90%, in the surroundings of the big cities of world [5]. The toxicity of DEP-PM is exerted mainly due to incorporation of polyaromatic hydrocarbons (PAHs). These DEP-PM are in nanometer in size and are penetrated very deep in the lung mucosa. Baecause of hydrophobic nature of PAH the DEP-PM can easily cross plasma memberane of cells and reach to the cytosol. Inside the cytosol it trigerrs many signalling mechanisms responsible for cell growth and differentiation [3]. It was observed that, lung toxicity increased in mainly childerun and they are at increased risk from diesel exhaust particles as compared to other as groups. Notably, there is transgenerational transmission of asthma risk if environmental particles are exposed in the pregnancy period [6]. The clinical manifestations due to DEPs exposure may include alteration of the respiratory functions, irritation of nose and eyes, fatigue and nausea and/or headache. The prolonged exposure of DEPs may produce respiratory function impairment along with cough and sputum [7]. After decades of research on the mechanism of DEPs induced toxicity, a majority of evidences support that, there is involvements of immune system and its mediators however the exact role is still not clearly understood [3, 8].

Mechanistically, it is understood that, remodelling of airway tissue is triggered by inflammatory mediaters released by different immune cells [9, 10]. In human asthma, the immunological mechanism responsible for the developments of initiation and progression of airway tisuue remodelling is still debated [11]. To understand the mechanism of tissue remodelling in asthma researchers use animal models, as theyare only available for remodelling in allergic asthma [9, 12]. In this line there are many relevant studies which highlight the important role of immune cells and their mediators in the development of airway tissue remodelling and subsequent fibrosis. Eosinophils are the main cell type and also play a key role in initiation of tissue remodelling of various tissues in many disease [13, 14]. It was very surprising to note that, obese mice when exposed to DEP are resistant and do not respond to eosinophilic airway inflammation [15]. Diesel exhaust particles showed stimulation of natural killer cells which in turn caused tissue remodelling and fibrosis [16]. It has recently been discovered that IL-23 paly a very important role in the initiation and development eosinophils mediated tissue remodelling and fibrosis [17]. IL-33 has also shown a very critical role in the development of diesel exhaust particles induced asthma pathophysiology [18]. We and others in earlier studies showed the role of interleukin (IL)-13 as a key mediator of airway tissue remodelling and fibrosis [19-21]. IL-13 is known to be pleiotropic nature of cytokine which is coded by a gene located on chromosome 5 [19]. It was observed that, IL-13 exerts its effect by triggering TGF production by varios immune cells including macrophages. In addition to airway remodelling, IL-13 also critically controlled the pathogenesis of nodular sclerosing Hodgkin’s disease, hepatic fibrosis, pulmonary fibrosis and progressive systemic sclerosis. Inspite the exact mechanism of IL-13 triggered tissue remodelling fibrosis has not yet been understood in details. There is still a lot of confusion in the mechanism of IL-13 mediated development of tissue remodelling and fibrosis.

The growing evidences highlights the role of eosinophils granules and fibroblast cells which are derived from immune cells [22, 23]. In a mice model of asthma, it was shown that allergic protein of soybean enhances the eosinophilic inflammation and exacerbation of asthmatic pathophysiology [24]. Eosinophil peroxidase was shown to oxidize the isoniazid and convert them in isoniazid-NAD active metabolite against M. tuberculosis [25]. Eosinophil peroxidase is also a one of the markers for atopy and bronchial hyper-responsiveness [26]. Eosinophil peroxidase helps in protein nitration during allergic airway inflammation in a murine model [27]. It is now being accepted that, lots of stem cells or immune cells comes to the lungs during any injury or allergen challenge and there differentiate into FSP-1 positive fibroblasts cells [22, 28]. How these fibroblast cells come to the lungs and affects tissue remodelling and fibrosis is still a big question and need extensive investigation. Despite the lots of research on the role of IL-13 and eosinophils in diesel exhaust particle induced tissue remodelling and fibrosis the exact mechanism associated to eosinophils role in it, is not properly known therefore the present study is planned to understand the role of eosinophils mediated tissue remodelling and fibrosis in the pathology of asthma.

## 2. Materials and Methods

### 2.1. Reagents and materials

Diesel exhaust particles (DEP) were generated and collected as per the protocol described earlier. Primary rabbit polyclonal antibodies against alpha-smooth muscle actin (a-SMA), collagen-1, eosinophil peroxidase (EPX) and Fibroblast specific protein-1 (FSP-1) were procured from Sigma-Aldrich (Merk). All other important reagents were procured from sigma other than specified Sigma-Aldrich (Merk).

### 2.2. Cell lines used

Human HL60 cells and A549 cell lines were used in the present study. Intitially these cells were procured from National Centre for Cell Sciences, Pune, India. After their purchage these cells are being continuously maintained at the tissue culture laboratory of ICMR-Vector Control Research Center, Puducherry.

### 2.3. Culture and passaging of the cells

DMEM or DMEM-ham’s F12 mixture of neutrients was used for culturing the 4549 cells. RPMI1640 was used to culture the HL-60 cells. In these mediums, 10% inactivated fetal bovine serum was supplemented. Cells were conditioned at 37°C in a 5% CO_2_/95% air and in a humidified atmosphere. After the adjustments of pH to 7.4 which was filter sterilized by filters of pore size of 0.22 μm.

### 2.4. Differentiation of HL-60 into eosinophils like cells

The HL-60 cell line used in these experiments was procured from NCCS, Pune, India and since then it has continuously been maintained in our tissue culture laboratory. The HL-60 cells before their differentiation were growing very well with fidelity. Terminal differentiation of HL-60 cell was induced by exposing the cells with 0.5 mM butyric acid for nearly 7 days. After the differentiation the exposure of various concentrations of soot or diesel exhaust particles were given and subsequent measurements of parameters were accomplished.

### 2.5. MTT assay for the assessment of cell viability or cell proliferation

MTT assay was perfoemed for the measurement of cell proliferation or cell viability [29]. Cells at the density of 10,000/well in 100 ul of volume were seeded in 96-well plate. After treatments the MTT reagent 3-(4,5-dimethylthiazol-2-yl)-2,5-diphenyltetrazolium bromide was suplimented to the wells (20 μl/well containing 100 μl of cell suspension). Color development was measured at 530 nm, using microplate reader.

### 2.6. Animal studies

BALB/c mice (8 to 12 weeks old) were used in this study. Before DEP intranasal challenge, the mice were anesthesized and 100 ug of DEP in 50 ul of saline were givin to the mice. After the intranasal chalage the mice organs were harvested and ket in the formalin and latter proccsed for immunohistochemistry

### 2.7. Immunohistochemistry of the tissue sections

The tissues of mice after collections were submerged in neutral buffered formalin for fixing. The tissues were embedded in paraffin and sectioned (5 µM thin). Sections were adhered to slides having positive charge. The 0.3% hydrogen peroxide in methanol was applied to quench endogenous peroxidase. Blocking with 1% BSA was done to block nonspecific sites. Tissue sections were incubated by primary antibodies at 4°C at 1:100 dilutions for a period of 12 hours. At 1:200 dilutions of HRP-conjugated secondary antibodies for a period of 1 hour was given at room temperature. Secondary antibodies alone were considered negative controls without the primary antibody. DAB-kit was used to further development of thse slides and counterstained using hematoxylin.

### 2.8. Isolation and quantification of RNA from lung tissue of mice

The lung sections of mice were lysed in 1000 µl trisol (crused) by pipetting up and down repeatedly (following the protocol of manufacture’s instructions sigma). The isolated RNA was kept for drying for 10 minutes. This dried RNA pellet was resdissolved in 20 µl of autoclaved purified water. The quantification of RNA was accomplished using Thermo Scientific µDrop™ Plate (Catalogue Number: N12391).

### 2.9. cDNA synthesis and real time PCR or qPCR for the expression analysis of inflammatory genes

The 200 ng/µl of the total isolated RNA was utilised for the construction of cDNA synthesis. Real time PCR or qPCR was performed to measure the relative expression of the inflammatory genes as listed in the Table No.1 due to exposure with DEP (diesel exhaust particles) in mice lungs. In brief, 5 µl of template (cDNA synthesized) was added to the reaction mixture containing 1 µl (0.5 picomoles) of gene specific primers, 3 µl of MiliQ water and 10 µl of SYBR Green. SYBR Green is a fluorescence dye and it selectively conjugate with dsDNA emitting fluorescence. The samples were assessed in duplicates. A negative control reaction was also put in which cDNA template was replaced with purified water. The C_T_ value obtained after the amplification of the inflammatory gene of interest was normalized with the reference gene (actin). Analysis of gene expression was accomplished using (The 2^-ΔΔCT^method). Results are expression as relative mRNA values of the each gene of interest.

### 2.10. Statistical analysis

Majority of results are expressed as mean ± S.E.M. Analysis of statistical procedures were achived by using one-way analysis of variance (ANOVA), followed by Tukey’s test. Graph-Pad prism 5 software was adopted for the statistical analysis. p value <0.05 was accepted as statistically significant.

## 3. Results

### 3.1. Effect of Diesel exhaust particles (DEP) on the cell proliferation of HL60 cells differentiated to eosinophils

Firstly, we have assessed the effects of DEP on HL-60 Cells. Prior to start the mechanistic effects of desel exhaust particles we have measured the cell viability of Hl-60 cells in response to DEP. As demonstrated by the bright field images in Fig. 1 A, DEP did not cause death of eosinophils cells indicating cell proliferation or cell survival. Fig. 1B demonstrated, DEP exposure did not cause cell death of these cells. DEP concentrations (125, 250, 500 and 1000 µg/ml) significantly increased the cell viability of HL-60 cells in dose dependent manner. As shown in Fig 1B. DEP did not cause death of these cells. As we have seen that DEP 500 ug/ml and 1000 ug/ml have significantly induced cell proliferation of HL-60 cells, differentiated to eosinophils. This phenomenon of DEP induced cell proliferation of eosinophil cells resemble as a default stage of the concerns tissue remodelling and fibrosis.

**Figure 1.**
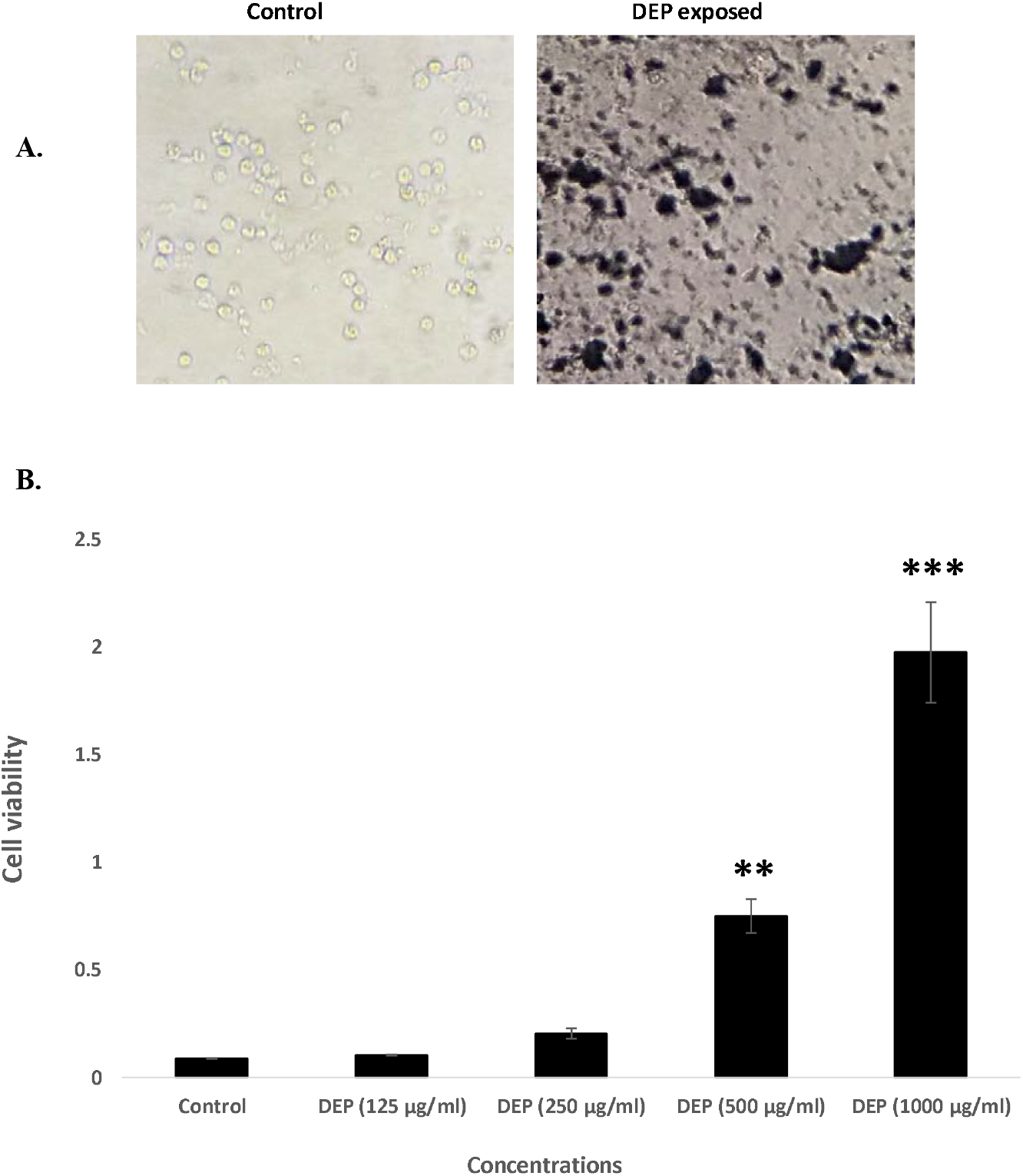
Dose dependent study of DEP on the cell proliferation of HL60 eosinophils cells. **A**. brightfield images of eosinophils in control and DEP exposed groups. **B**. Histograms are showing the dose dependent study of DEP on the cell proliferation (measured by MTT) of HL-60 eosinophil cells. ** p<0.01 compared and *** p<0.001 compared with control.

### 3.2. Effect of Etoricoxib, on Disesal Exhaust Particle triggered release of IL-13 by the HL-60 cells, differentiated to eosinophils

We have further tested that, whether “Etoricoxib”, a selective cox-2 inhibiter, inhibit diesel exhaust particles (DEP) induced IL-13 release or not in eosinophils cells. To accomplish this HL-60, eosinophil cells were treated with “Etoricoxib” concentrations (1 ug/ml and 2 ug/ml) prior to DEP exposure. After this, HL-60 cells were exposed with the DEP (500 ug/ml) and measurement of IL-13 were done in the cell supernatant collected after 24 hours of treatments. As shown in the Fig. 2, Etoricoxib does not inhibit DEP induced Interleukin-13 release. It also shows that COX-2 inhibition is not linked with the IL-13 pathway and DEP follows a different pathway which may be independent of IL-13.

**Figure 2.**
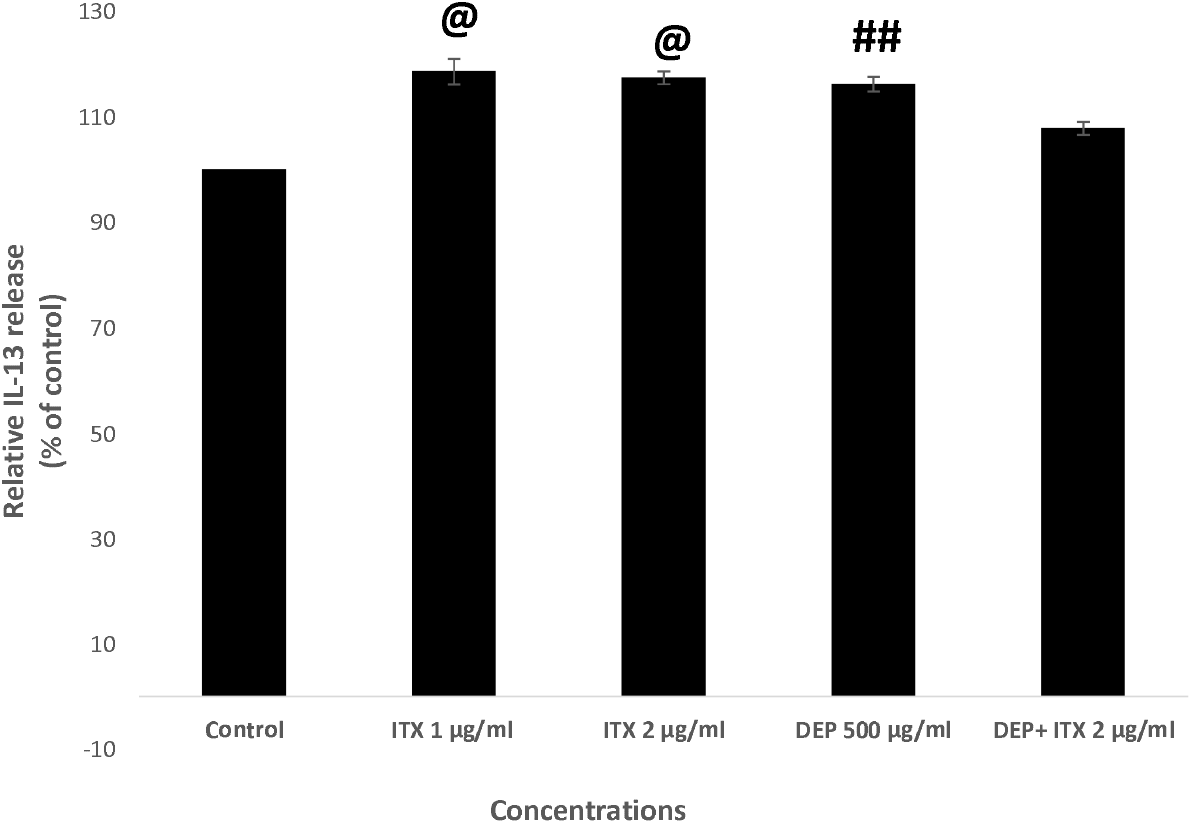
Effects of Etoricoxib on the DEP induced IL-13 release in HL-60 eosinophils cells. Histograms represent the cell IL-13 release by HL-60 differentiated eosinophils and its inhibition by Etoricoxib. @ p<0.05 compared with control, ## p<0.01 compared with control.

### 3.3. The secretome from DEP-induced eosinophils restrict cell proliferation of lung, A549 epithelial cells

It has been noticed that the eosinophils effects lung epithelial and other cells in a variety of ways and may be affecting cell proliferation or cell death. Therefore, we have tested the effect of secretome obtained from DEP-induced eosinophils on the cell proliferation and apoptosis of lung epithelial cells. DEP-induced the release of different cell mediators in the medium which may be producing its effect on the cell proliferation or induction of cell death. As seen in the Fig. 3. we found that diesel exhaust particle-induced proliferation of A549 cells, is significantly inhibited by the secretome obtained from eosinophils. Secretome from 250 µg/ml and 500 µg/ml treated cells has significantly decreased the cell proliferation of A549 cells in a 24 hours duration. On the other had DEP has significantly increased the number of the A549 cells. This finding has confirmed that DEP induces eosinophils to release some mediators of inflammation which controls the pathological changes in the lungs tissue and restrict tissue remodelling.

**Figure 3.**
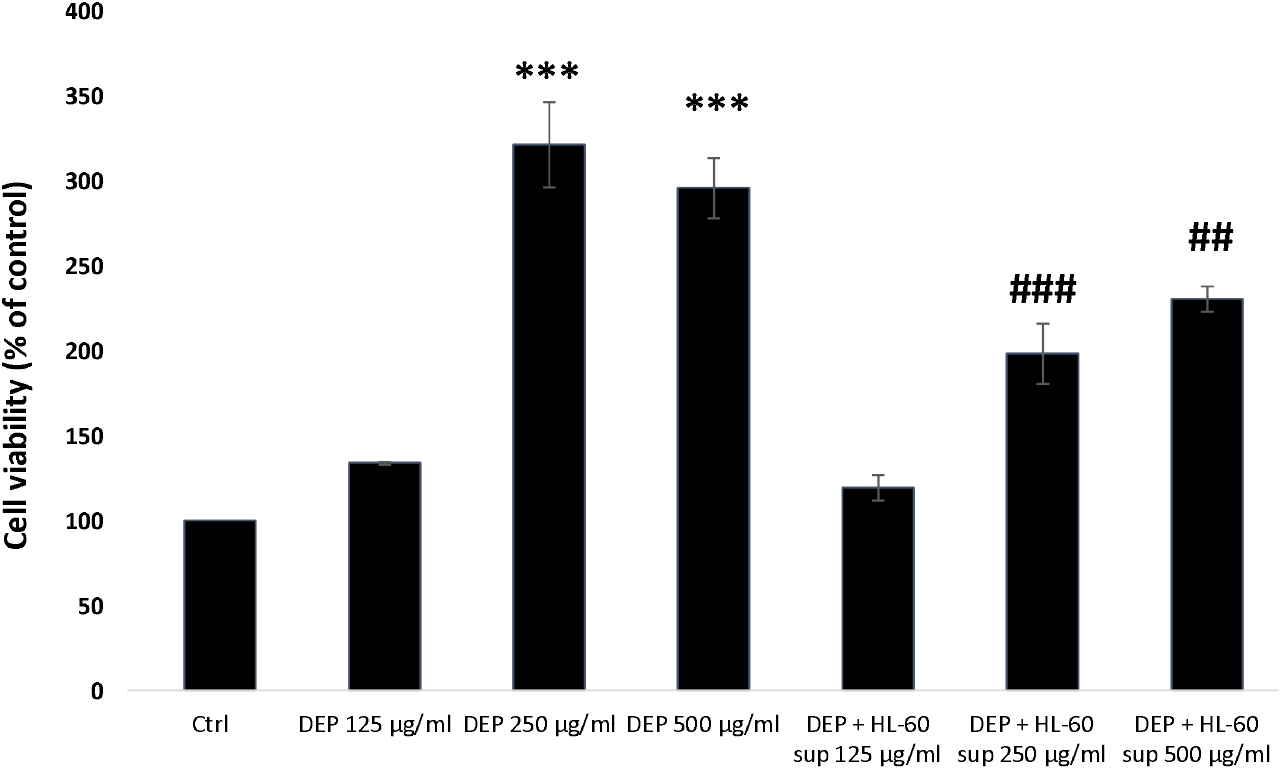
Effects of eosinophils secretome on the DEP-induced cell proliferation of lung epithelial A549 cells. Histograms represent the cell viability (measured by MTT assay) of the A549 cells in the presence and absence of secretome collected from DEP-exposed eosinophils. *** p<0.001 compared with control, ## p<0.01 and ### p<0.001 compared with DEP treated group.

### 3.4. Purified Interleukin-13 has inhibited cell proliferation of A549 cells

Further, we speculated that, IL-13 released by eosinophils may be a limiting factor and may restrict epithelial cell proliferation. To test this hypothesis, we have exposed lung epithelial cells with different concentrations of purified IL-13 (0.001, 0.01 and 01 ng/ml) for 24 hours and measured cell viability by MTT assay. As seen in the Fig. No. 4. interleukin-13 has significantly inhibited the cell proliferation of A549 in dose dependent manner. This confirmed that, eosinophils release Interleukin-13 in response to DEP exposure to control the cell proliferation of lung epithelial cells which in turn are involved in the development of tissue remodelling and fibrosis.

**Figure 4.**
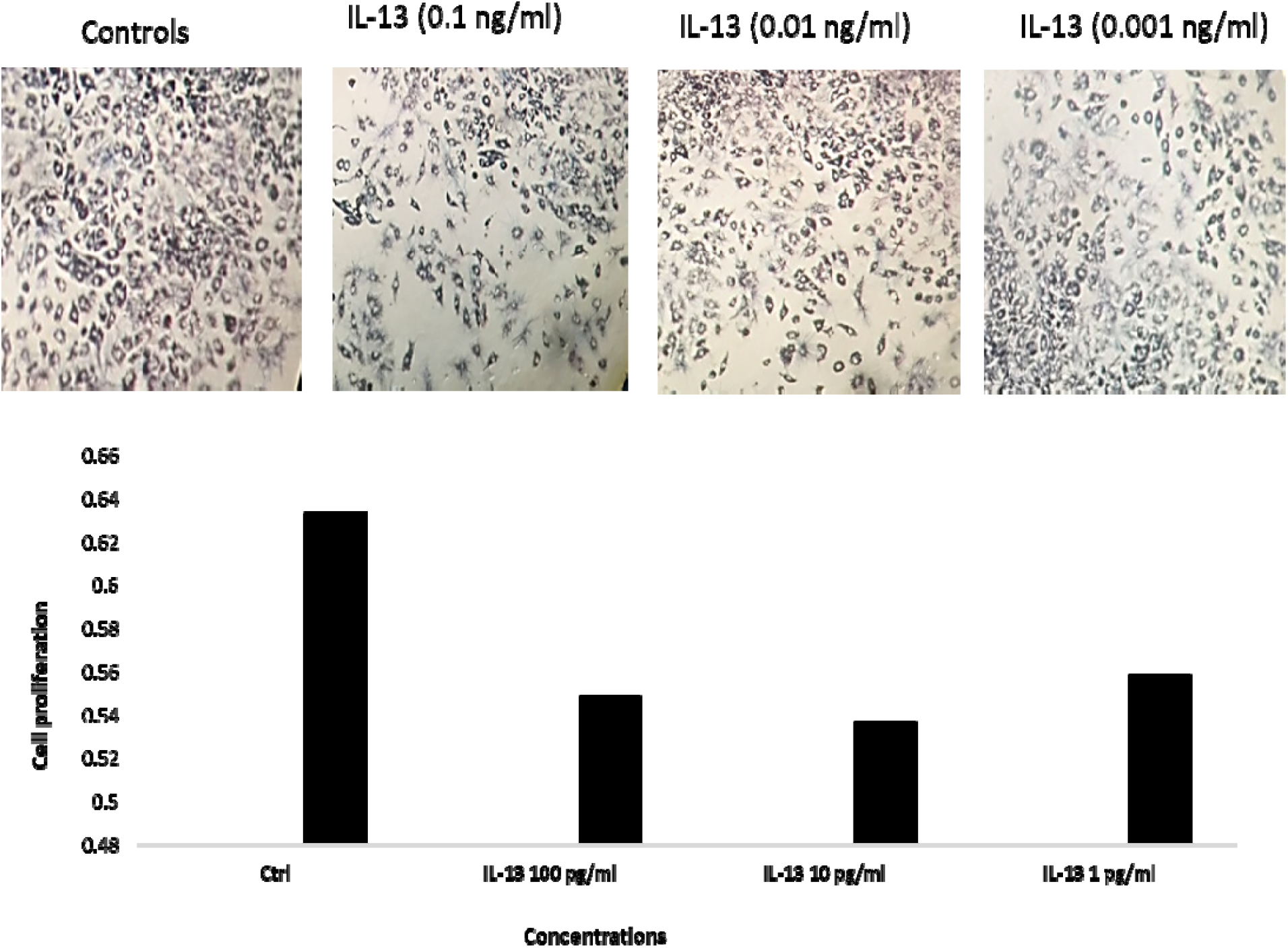
Effects of purified IL-13 on the cell proliferation of lung epithelial A549 cells. Images shows different concentrations of IL-13 exposure to the A549 cells. MTT assay was conducted to assess the cell proliferation of A549 cells and after that images were captured of A549 cells exposed with different concentrations of interleukin-13. Histograms represent the cell viability of the A549 cells in the presence and absence of secretome collected from DEP exposed eosinophils.

### 3.5. Effect of Diesel exhaust particles (DEP) on eosinophil recruitment and degranulation, fibroblast proliferation in mice lungs in vivo

Next, we have measured eosinophils infiltration and recruitment to the lungs and their degranulation in response to dep exposure to the mice. To accomplish that, we have measured eosinophil peroxidase (EPX) expression and localization in the mice lungs using immune-histochemistry. As shown in the Fig. No. 5, DEP exposure to the mice lungs increase the EPX immunostaining compared to saline challenged mice. This EPX staining directly correlate to the eosinophil’s recruitments and degranulation due to DEP exposure in mice lungs. It is further interesting to note that eosinophil peroxidase is an important factor which will be upregulated in the development of tissue remodelling and fibrosis.

**Figure 5.**
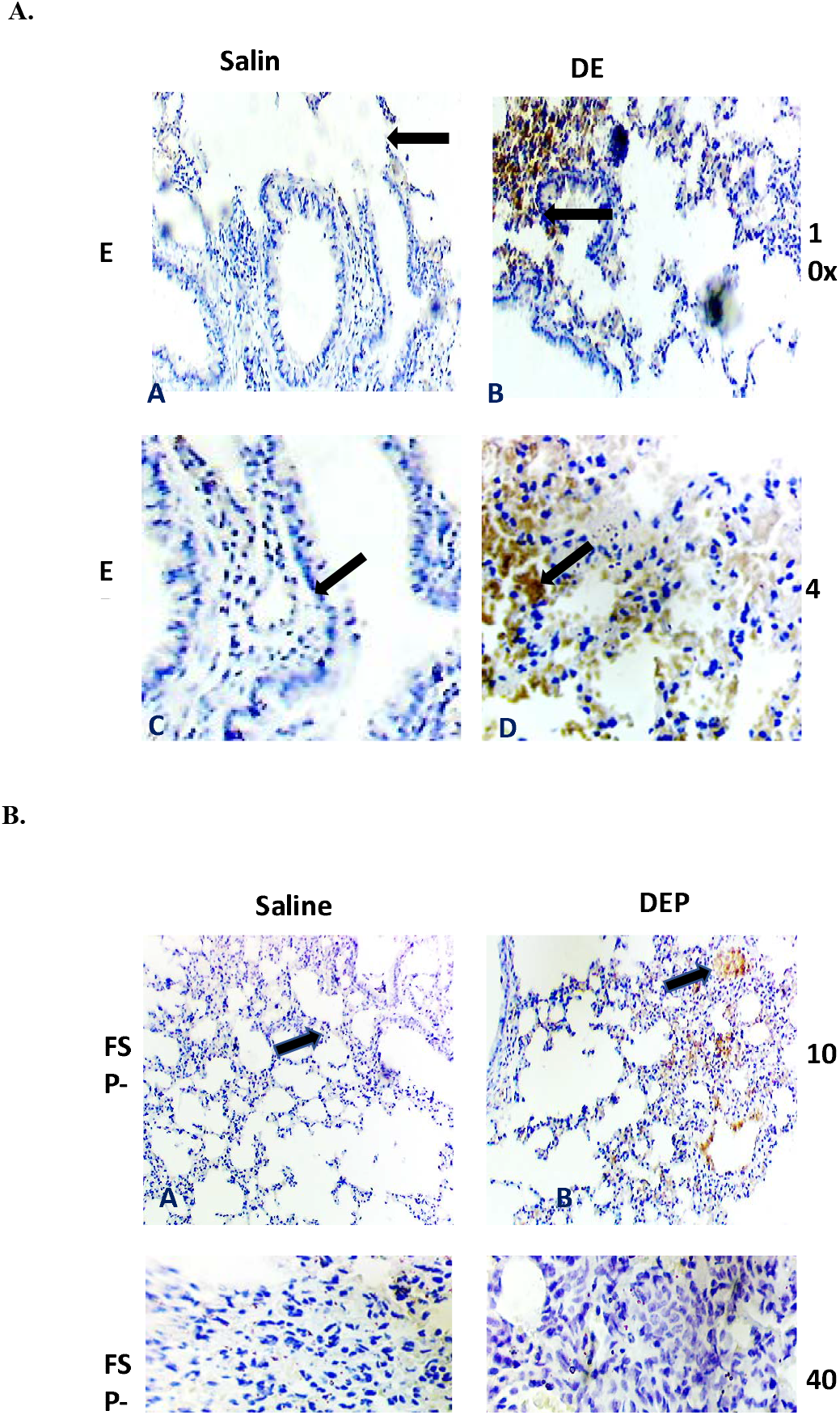
Effects of DEP on the Eosinophil peroxidase (EPX) and fibroblast specific protein-1 (FSP-1) in mice lung. A. Images in the figure show expression and localization prolife EPX in saline and diesel exhaust particles treated mice. B. Images in the figure show expression and localization of FSP-1 protein in saline and diesel exhaust particles treated mice.

Similarly, we have measured fibroblast recruitments or conversion in mice lungs in response to DEP exposure. To achieve this, we have measured fibroblast specific protein-1 (FSP-1) expression and localization in the mice lungs using immune-histochemistry. As shown in the Fig. No. 5B, DEP exposure to the mice lungs increase the FSP-1 immunostaining compared to saline challenged mice. The presence of FSP-1 staining indicate, the presence of fibroblast which in turn secrete a lot of fibrous proteins involved in the initiation and progression of tissue remodelling and fibrosis.

### 3.6. Effect of Diesel exhaust particles (DEP) on alpha SMA expressions and localization in mice lungs in vivo

We have further measured alpha SMA expression in order to measure the tissue remodelling and fibrosis in the mice lungs. As shown in the Fig. No. 6, DEP exposure to the mice lungs increased the alpha SMA immunostaining compared to saline challenged mice. Similarly, we have measured tissue remodelling bu H&E staining. As shown in the Fig. No. 6A, DEP exposure to the mice lungs increase the cell numbers compared to saline challenged mice.

**Figure 6.**
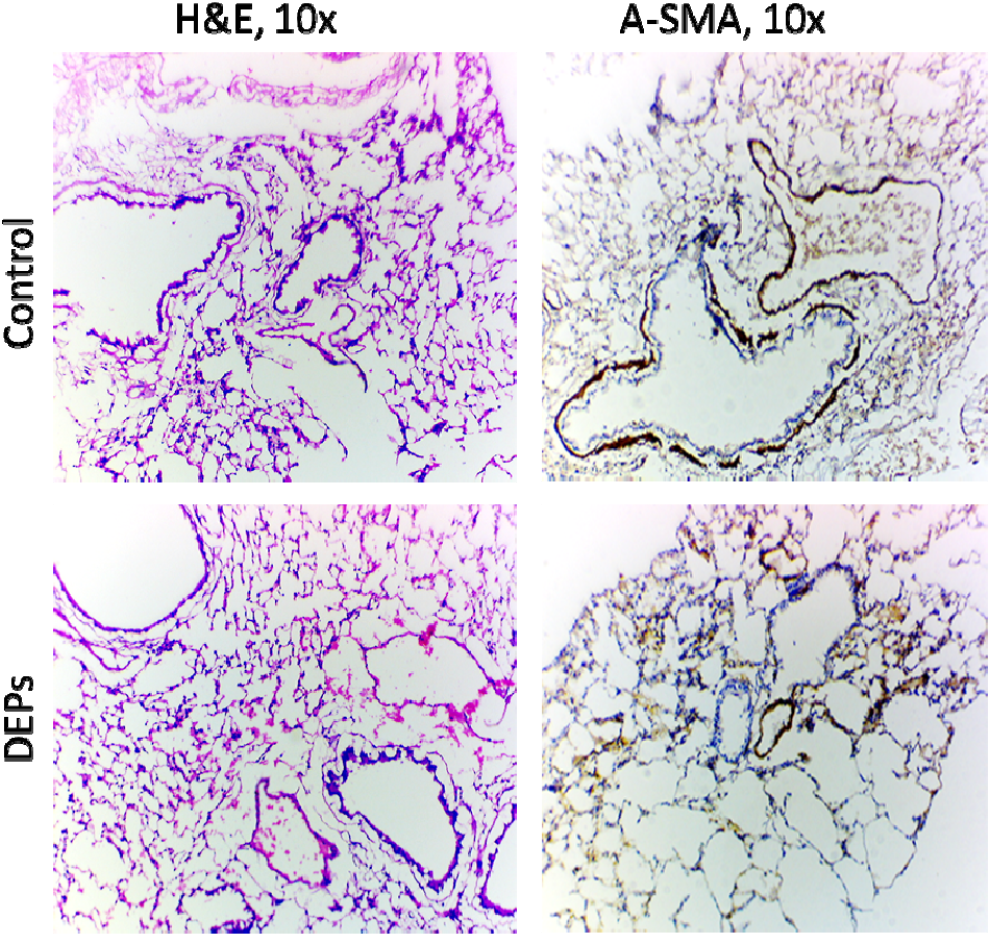
Effects of DEP on the alpha-SMA expression and localozation in mice lung. A. Images in the figure show expression and localization prolife EPX in saline and diesel exhaust particles treated mice. B. Images in the figure show expression and localization of FSP-1 protein in saline and diesel exhaust particles treated mice.

### 3.7. Effect of diesel exhaust particles (DEP) on mRNA expressions analysis of inflammatory cytokines gene in mice lungs in vivo

Mediaters of inflammation plays an important role in the initiation of progression of tissue remodelling and fibrosis. Therefore, we have asseded the mRNA expressions of inflammatory cytokine genes, i.e. IL-13, IFN-Y, TNF-beta, IL-4, IL-5, MBP and FcεRIα in saline and DEP treated mice lungs. mRNA expression was measured using qPCR as per the protocol mentioned in the material and method sections and in the table No.1. As mentioned in the Fig. No. 7, DEP exposure to the mice lungs has incresed the mRNA expression of, i.e. IL-13, IFN-Y, IL-4, IL-5 and FcεRIα. However it has decresed the expression of MBP (major basic protein) and TNF-beta mRNA expression. These expression anlysis significantly shows that there is changes in the cytokine level in the lungs of mice due to DEP.

**Table No. 1.** Details of primers used in the study

| Sl No | Primers | Annealing Temp | Amplicon Size |
| --- | --- | --- | --- |
| 1. | IFNgama F - 5' gCT CTg AgA CAA TgA ACg CT 3' (20 mer) | 56°C | 229bp |
|  | IFNgama R - 5' AAA gAg ATA ATC Tgg CTC TgC (21 mer) |  |  |
| 2 | TNF Beta F - 5' ACC gCA ACA ACg CCA TCT AT (20 mer) | 57.9°C | 200 bp |
|  | TNF Beta R - 5' gTA ACg CCA ggA ATT gTT gC (20 mer) |  |  |
| 3. | IL 4 -F- 5' TCg gCA TTT TgA ACg Agg TC (20 mer) | 56°C | 216bp |
|  | IL 4 -R - 5' gAA AAg CCC gAA AgA gTC TC (20 mer) |  |  |
| 4. | IL 5 -F - 5' TCA CCg AgC TCT gTT gAC AA (20 mer) | 60°C | 201 bp |
|  | IL 5 -R - 5' CCA CAC TTC TCT TTT Tgg Cg (20 mer) |  |  |
| 5. | IL 13 -F - 5' gCC AgC CCA CAg TTC TA CAg C (22 mer) | 60°C | 182bp |
|  | IL 13 -R - 5' gTg ATg TTg CTC AgC TCC TCA (21 mer) |  |  |
| 6. | FcεRIα F - 5' TgC CCA gCT gTg CCT AgC AC (20 mer) | 58°C | 527bp |
|  | FcεRIα R - 5' CCA CCT gCC TAA gAT AgC CC (20 mer) |  |  |
| 7. | MBP F - 5' gAg CgT CTg CTC TTC ATC TgA (21 mer) | 56.1°C | 1282bp |
|  | MBP R - 5' CAC TgA AAC TgT gAA Tgg Agg C (22 mer) |  |  |

**Figure 7.**
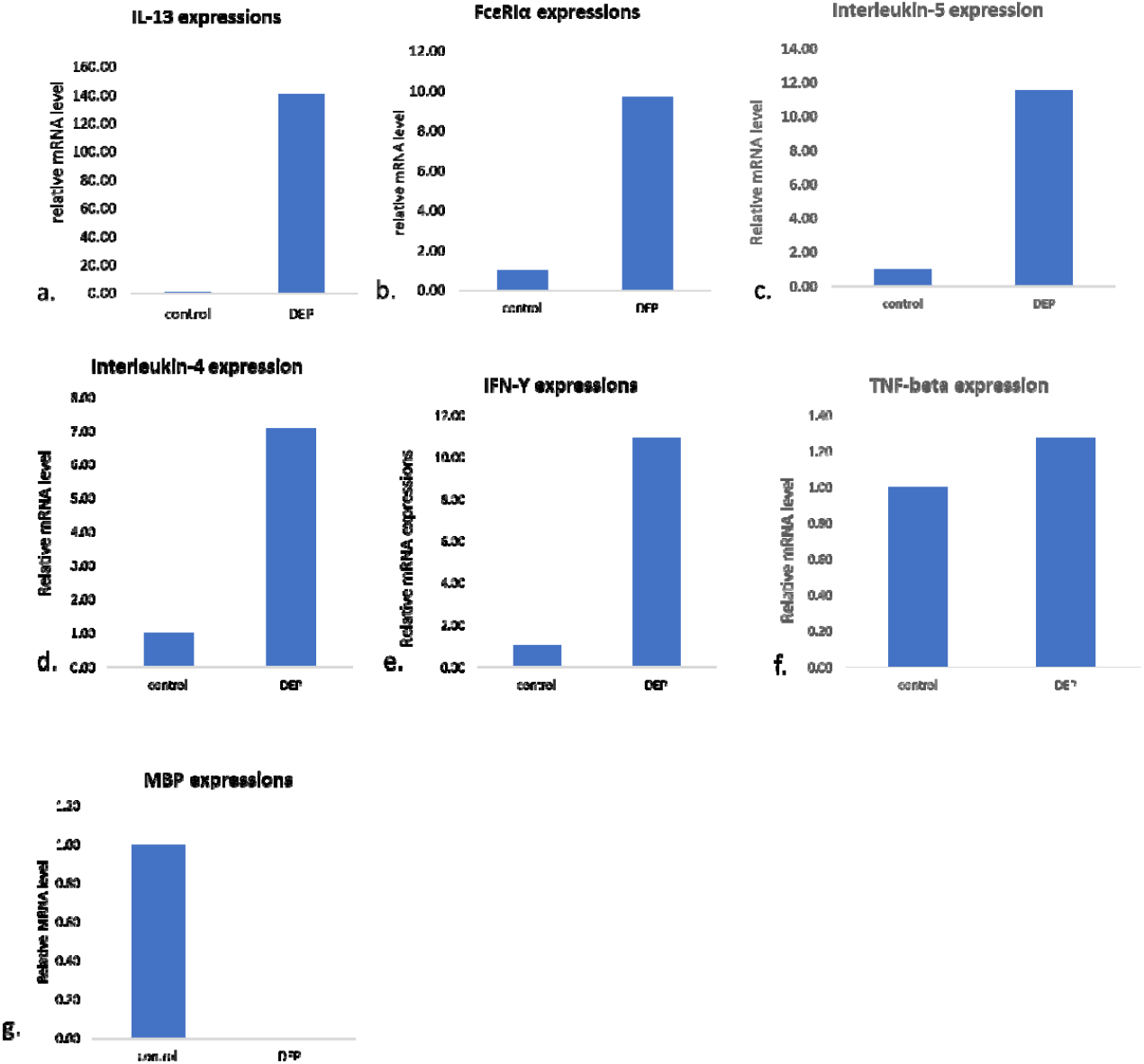
The mRNA expression profile inflammater cytokine genes in mice lungs after DEP exposure using Real time PCR. **A**. Real time PCR measured expression profile to IL-13, B. FcεRIα, C. IL-4, D. IL-5, E. MBP, F. IFN-Y, G. TNF-beta. Beta action was taken as an internal control. mRNA level is expressed as relative mRNA expression of specific cytokine genes.

## 4. Discussion

In the present study, we show that there is clear-cut involvement of interleukin-13 in diesel exhaust particle (DEP) induced tissue remodelling and fibrosis [4]. We show that, interleukin-13, restrict cell proliferation of lung epithelial cells [30]. We also show that, eosinophils participate in the development of tissue remodelling by secreting interleukin-13 which in turn restrict tissue remodelling and fibrosis.

We know that eosinophils play very important role in the development of tissue remodelling and fibrosis [13, 31]. It is father important to note that various mediators of inflammation are upregulated in the development of tissue remodelling and fibrosis [32, 33]. The interactions of immune cells specially eosinophils to the lung epithelial cells is an important mechanism for the development of tissue remodelling and fibrosis [10]. Recently we have elucidated that there are two distinct type of eosinophils subsets that regulate the initiation and progression of pathological process in various diseases [34]. We have also elucidated that IL-15 induce a huge number of tissue eosinophils in lungs and produce a protective effect. In hour and other research, it was known that eosinophils may me of two district type and functions in a certain direction based on the various immune regulators [35, 36]. In the present study, we have shown that, eosinophils respond to the diesel exhaust particles by releasing interleukin-13 and other mediators which restrict the epithelial cell proliferation however what is the role of IL-13 on other cell types of lungs is not properly known.

Interleuki-13 has widely been understood to cause or promote the development of tissue remodelling and fibrosis [37, 38]. It has already been known that, diesel exhaust particles induce development of lung tissue pathophysiology and tissue remodelling, however its mechanism is still largely unknown [2, 4]. In a previous study, it was also shown that, tobacco smoke induced IL-13 release is involved the structural changes in the lungs [39]. ERK1/2 mitogen-activated protein kinase selectively mediates IL-13-induced lung inflammation and remodelling in vivo [40]. Further it is also understood that there are therapeutic effects of IL-13 immunoneutralization during chronic experimental fungal asthma, suggesting an important role in the pathogenesis of asthma [41]. In the present study, it is proved that eosinophils secrete some mediators of inflammation which are responsible for the inhibition of cell proliferation [42]. It is also further important to note that, this mechanism of tissue remodelling and fibrosis mediated by the interleukin-13 is an important fact which need to be understood in details [33]. We have previously published that allergen induced tissue remodelling and fibrosis is independent of interleukin-13 [21]. That means allergen can induce tissue remodelling and fibrosis in absence of interleukin-13. This finding suggests that there may be other role of interleukin-13 other than inducing tissue remodelling and fibrosis in the disease. In the present study, we show that IL-13 when induced by allergen (in this study DEP), shows a restricted on the induced cell proliferation of lung epithelial cells in vitro. This finding is very important towards the important role of interleukin-13 in limiting cell proliferation of lung cells. There are also many studies in which interleukin-13 has shown its protective role including neuroimmune functions [43-45].

Interleukin-13 and IL-4 are known to supresses the COX-2 signalling in a variety of studies but how this is linked it is not at clearly known [46, 47]. It was described that, IL-4 and IL-13 suppress the release of prostaglandins by follicular dendritic cells by repressing expression profile of COX-2 via JAK1 and STAT6 pathway [48]. COX-2 inhibition is also linked to cell proliferation and release of various mediators of inflammation both *in vivo* and *in vitro*. Therefore, we have used “Etoricoxib” a selective inhibitor of COX-2 enzyme, to understated the releases of inter interleukin-13 by DEP exposed eosinophil cells. Contrary to other study, we found that Etoricoxib does not significantly down regulate the IL-13 release by DEP-exposed eosinophils indicating a different intracellular mechanism of action. The findings in the present study are also similar and IL-13 release is not inhibited by the COX-2 inhibiter Etoricoxib, providing a support to the authenticity and validity of role of IL-13 in this study.

It is very interesting to know that eosinophils granules mainly eosinophil peroxidase (EPX) produce very important role in the development of tissue remodelling and fibrosis [23, 49, 50]. Eosinophil peroxidase is also one of the key marker proteins being used as the detection of eosinophils in tissues [51]. In an interesting study it was shown that mice knockout with eosinophil peroxidase (EPX) significantly develop lung pathology indicating an important role of eosinophils in the attenuation of lung tissue remodelling or pathophysiological conditions [52]. On the other hand, Fibroblast specific protein-1 is believed to be a string marker for fibroblast [53]. Its expression positivity shows the presence of fibroblasts in any tissue. We have in our recent study has published that FSP-1 positive cells appear in the lungs during the tissue remodelling and fibrosis and significantly decrease whenever tissue remodelling is reduced or attenuated [54]. It is interesting to note that, some of the monocytes can be converted in the fibroblasts and behave like fibroblasts [22]. These cells are called monocytes-derived fibroblasts [22]. It is speculated that, DEP induced monocytes to come and differentiate into fibroblasts like cells and then they in turn starts producing collagen and fibrous proteins responsible for tissue fibrosis [22, 55]. In the present study EPX and fibroblast specific protein-1 (FSP-1) is also are also upregulated in the mice lungs exposed to DEP [51, 56]. While EPX shows the important role of eosinophils degranulation and infiltration to the lungs. FSP-1 positive cells show the recruitment of fibroblasts which in turn secrete lot of fibrous proteins such as collagen, fibronectin tenascin and others which constitute fibrosis in the lung tissue. It is further important to know that, role of fibroblast specific protein-1 is still need not be understood in details during the development of tissue remodelling and fibrosis [57].

In conclusion it can be said that, interleukin-13 is upregulated and released by the eosinophils cells which are recruited in response to environmental allergens (DEP) to restrict the tissue remodelling and fibrosis. Interleukin-13 treatment reduces the cell proliferation of epithelial which depicts that, eosinophils mediated IL-13 release may be a beneficial event in controlling the tissue remodelling and fibrosis in asthma pathophysiology. The present findings are encouraging however the further research is needed to reclarify the things.

## Acknowledgements

The “young scientist grant” to Dr. Rituraj Niranjan grant No. SB/YS/LS-198/2014 by SERB (DST, India.) is acknowledged gratefully.

## Conflict of interest

None.

### Abbreviations

AD: aerodynamic diameter
COX-2: cyclooxygenase-2
HL-60: Human Leukemia Cell Line 60
HRP: Horse Radis Peroxidase
EPX: Eosinophil peroxidase
α-SMA: alpha-smooth muscle actin
FSP-1: fibroblast specific protein
DEP: Diesel exhaust particles
IL-13: interleukin-13
ROS: Reactive oxygen species
HBSS: Hank’s balanced salt solution
Th: T helper cell
Ig: Immunoglobulin
PM: particulate matter
TGF-β: Tumour growth factor-beta
PAHs: polycyclic aromatic hydrocarbons
IFN-γ: interferon gamma
TNF-β: tumour necrosis facter beta
IL-4: Interleukin-4
IL-5: interleukin-5
MBP: major basic protein
FcεRIα: Fc epsilon RecepterI alpha.

---

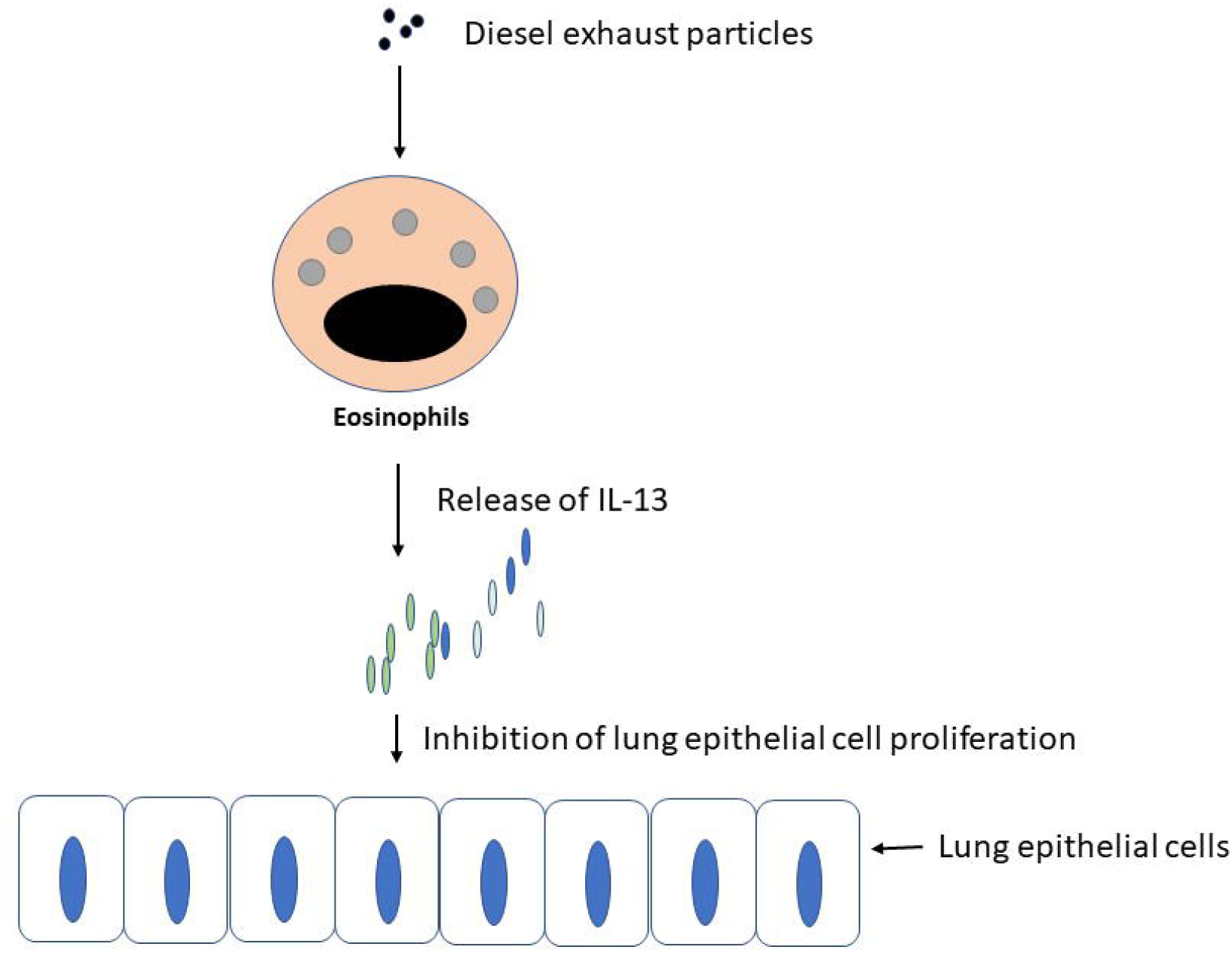

## HIGHLIGHTS OF THE ARTICLE

1. We found, diesel exhaust particles (DEP) induce eosinophils proliferation and interleukin-13 release *in vitro*.

2. Eosinophils cells significantly restrict cell proliferation of epithelial cells induced by diesel exhaust particles.

3. Purified interleukin-13 decreases the proliferation of lung epithelial, A549 cells.

4. Etoricoxib, (selective COX-2 inhibitor) did not inhibit DEP-triggered release of interleukin-13 by eosinophils.

5. Altered expressions of IL-13, FcεRIα, IL-5, IL-4, IFNgama, TNF-beta genes were found by eosinophils due to DEP.

## References

1. Baulig A, Blanchet S, Rumelhard M, Lacroix G, Marano F, Baeza-Squiban A. Fine urban atmospheric particulate matter modulates inflammatory gene and protein expression in human bronchial epithelial cells. Front Biosci.2007;12:771–782.

2. Liu J, et al. Diesel exhaust inhalation exposure induces pulmonary arterial hypertension in mice. Environ Pollut.2018;237:747–755.

3. Niranjan R, Thakur AK. The Toxicological Mechanisms of Environmental Soot (Black Carbon) and Carbon Black: Focus on Oxidative Stress and Inflammatory Pathways. Front Immunol.2017;8:763.

4. Kim BG, et al. Long-Term Effects of Diesel Exhaust Particles on Airway Inflammation and Remodeling in a Mouse Model. Allergy Asthma Immunol Res.2016;8:246–256.

5. Van Winkle LS, Brown CD, Shimizu JA, Gunderson AD, Evans MJ, Plopper CG. Impaired recovery from naphthalene-induced bronchiolar epithelial injury in mice exposed to aged and diluted sidestream cigarette smoke. Toxicol Lett.2004;154:1–9.

6. Gregory DJ, Kobzik L, Yang Z, McGuire CC, Fedulov AV. Transgenerational transmission of asthma risk after exposure to environmental particles during pregnancy. Am J Physiol Lung Cell Mol Physiol.2017;313:L395–L405.

7. Peden DB, Bush RK. Advances in environmental and occupational disorders in 2013. J Allergy Clin Immunol.2014;133:1265–1269.

8. Alexis NE, Carlsten C. Interplay of air pollution and asthma immunopathogenesis: a focused review of diesel exhaust and ozone. Int Immunopharmacol.2014;23:347–355.

9. Inoue K, et al. Effects of naphthoquinone on airway responsiveness in the presence or absence of antigen in mice. Arch Toxicol.2007;81:575–581.

10. Amin K, Ekberg-Jansson A, Lofdahl CG, Venge P. Relationship between inflammatory cells and structural changes in the lungs of asymptomatic and never smokers: a biopsy study. Thorax.2003;58:135–142.

11. Wooding DJ, et al. Acute air pollution exposure alters neutrophils in never-smokers and at-risk humans. Eur Respir J.2020;55.

12. Van Den Broucke S, Vanoirbeek J, Alfaro-Moreno E, Hoet P. Contribution of mast cells in irritant-induced airway epithelial barrier impairment in vitro. Toxicol Ind Health.2020:748233720948771.

13. Vohralik EJ, Psaila AM, Knights AJ, Quinlan KGR. EoTHINophils: Eosinophils as key players in adipose tissue homeostasis. Clin Exp Pharmacol Physiol.2020;47:1495–1505.

14. Clark K, et al. Eosinophil degranulation in the allergic lung of mice primarily occurs in the airway lumen. J Leukoc Biol.2004;75:1001–1009.

15. Yanagisawa R, Koike E, Ichinose T, Takano H. Obese mice are resistant to eosinophilic airway inflammation induced by diesel exhaust particles. J Appl Toxicol.2014;34:688–694.

16. Qian Q, et al. Maternal diesel particle exposure promotes offspring asthma through NK cell-derived granzyme B. J Clin Invest.2020;130:4133–4151.

17. Lee HS, Park DE, Lee JW, Sohn KH, Cho SH, Park HW. Role of interleukin-23 in the development of nonallergic eosinophilic inflammation in a murine model of asthma. Exp Mol Med.2020;52:92–104.

18. Brandt EB, Bolcas PE, Ruff BP, Khurana Hershey GK. IL33 contributes to diesel pollution-mediated increase in experimental asthma severity. Allergy.2020.

19. Wang X, et al. IL-4/IL-13 upregulates Sonic hedgehog expression to induce allergic airway epithelial remodeling. Am J Physiol Lung Cell Mol Physiol.2020;318:L888–L899.

20. Upparahalli Venkateshaiah S, et al. Attenuation of Allergen-, IL-13-, and TGF-alpha-induced Lung Fibrosis after the Treatment of rIL-15 in Mice. Am J Respir Cell Mol Biol.2019;61:97–109.

21. Niranjan R, Rayapudi M, Mishra A, Dutt P, Dynda S, Mishra A. Pathogenesis of allergen-induced eosinophilic esophagitis is independent of interleukin (IL)-13. Immunol Cell Biol.2013;91:408–415.

22. Abe S, et al. Functional analysis of human fibrocytes derived from monocytes reveals their profibrotic phenotype through paracrine effects. J Med Invest.2020;67:102–112.

23. Acharya KR, Ackerman SJ. Eosinophil granule proteins: form and function. J Biol Chem.2014;289:17406–17415.

24. Alvarez-Simon D, Munoz X, Gomez-Olles S, de Homdedeu M, Untoria MD, Cruz MJ. Effects of diesel exhaust particle exposure on a murine model of asthma due to soybean. PLoS One.2017;12:e0179569.

25. Babu D, et al. Eosinophil peroxidase oxidizes isoniazid to form the active metabolite against M. tuberculosis, isoniazid-NAD(). Chem Biol Interact.2019;305:48–53.

26. Bjornsson E, Janson C, Hakansson L, Enander I, Venge P, Boman G. Eosinophil peroxidase: a new serum marker of atopy and bronchial hyper-responsiveness. Respir Med.1996;90:39–46.

27. Duguet A, Iijima H, Eum SY, Hamid Q, Eidelman DH. Eosinophil peroxidase mediates protein nitration in allergic airway inflammation in mice. Am J Respir Crit Care Med.2001;164:1119–1126.

28. Yan D, Liu X, Xu H, Guo SW. Platelets induce endothelial-mesenchymal transition and subsequent fibrogenesis in endometriosis. Reprod Biomed Online.2020.

29. Mosmann T. Rapid colorimetric assay for cellular growth and survival: application to proliferation and cytotoxicity assays. J Immunol Methods.1983;65:55–63.

30. Konstantinidis AK, et al. Cellular localization of interleukin 13 receptor alpha2 in human primary bronchial epithelial cells and fibroblasts. J Investig Allergol Clin Immunol.2008;18:174–180.

31. Niranjan R, et al. Involvement of interleukin-18 in the pathogenesis of human eosinophilic esophagitis. Clin Immunol.2015;157:103–113.

32. Han XP, et al. EPO modified MSCs can inhibit asthmatic airway remodeling in an animal model. J Cell Biochem.2018;119:1008–1016.

33. Lee MY, et al. Anti-inflammatory and anti-allergic effects of kefir in a mouse asthma model. Immunobiology.2007;212:647–654.

34. Venkateshaiah SU, et al. A critical role for IL-18 in transformation and maturation of naive eosinophils to pathogenic eosinophils. J Allergy Clin Immunol.2018;142:301–305.

35. Mavi P, et al. Allergen-induced resistin-like molecule-alpha promotes esophageal epithelial cell hyperplasia in eosinophilic esophagitis. Am J Physiol Gastrointest Liver Physiol.2014;307:G499–507.

36. Rayapudi M, et al. Invariant natural killer T-cell neutralization is a possible novel therapy for human eosinophilic esophagitis. Clin Transl Immunology.2014;3:e9.

37. Ma Y, et al. Sustained suppression of IL-13 by a vaccine attenuates airway inflammation and remodeling in mice. Am J Respir Cell Mol Biol.2013;48:540–549.

38. Zuo L, et al. IL-13 induces esophageal remodeling and gene expression by an eosinophil-independent, IL-13R alpha 2-inhibited pathway. J Immunol.2010;185:660–669.

39. Cooper PR, Poll CT, Barnes PJ, Sturton RG. Involvement of IL-13 in tobacco smoke-induced changes in the structure and function of rat intrapulmonary airways. Am J Respir Cell Mol Biol.2010;43:220–226.

40. Lee PJ, et al. ERK1/2 mitogen-activated protein kinase selectively mediates IL-13-induced lung inflammation and remodeling in vivo. J Clin Invest.2006;116:163–173.

41. Blease K, Jakubzick C, Westwick J, Lukacs N, Kunkel SL, Hogaboam CM. Therapeutic effect of IL-13 immunoneutralization during chronic experimental fungal asthma. J Immunol.2001;166:5219–5224.

42. Wadsworth SJ, Atsuta R, McIntyre JO, Hackett TL, Singhera GK, Dorscheid DR. IL-13 and TH2 cytokine exposure triggers matrix metalloproteinase 7-mediated Fas ligand cleavage from bronchial epithelial cells. J Allergy Clin Immunol.2010;126:366-374, 374 e361-368.

43. Mori S, Maher P, Conti B. Neuroimmunology of the Interleukins 13 and 4. Brain Sci.2016;6.

44. Offner H, et al. Treatment of passive experimental autoimmune encephalomyelitis in SJL mice with a recombinant TCR ligand induces IL-13 and prevents axonal injury. J Immunol.2005;175:4103–4111.

45. Koning N, et al. Expression of the inhibitory CD200 receptor is associated with alternative macrophage activation. J Innate Immun.2010;2:195–200.

46. Onoe Y, et al. IL-13 and IL-4 inhibit bone resorption by suppressing cyclooxygenase-2-dependent prostaglandin synthesis in osteoblasts. J Immunol.1996;156:758–764.

47. Saito A, Okazaki H, Sugawara I, Yamamoto K, Takizawa H. Potential action of IL-4 and IL-13 as fibrogenic factors on lung fibroblasts in vitro. Int Arch Allergy Immunol.2003;132:168–176.

48. Cho W, Kim Y, Jeoung DI, Kim YM, Choe J. IL-4 and IL-13 suppress prostaglandins production in human follicular dendritic cells by repressing COX-2 and mPGES-1 expression through JAK1 and STAT6. Mol Immunol.2011;48:966–972.

49. Wills-Karp M, Karp CL. Biomedicine. Eosinophils in asthma: remodeling a tangled tale. Science.2004;305:1726–1729.

50. Percopo CM, et al. Impact of eosinophil-peroxidase (EPX) deficiency on eosinophil structure and function in mouse airways. J Leukoc Biol.2019;105:151–161.

51. Wright BL, et al. Image Analysis of Eosinophil Peroxidase Immunohistochemistry for Diagnosis of Eosinophilic Esophagitis. Dig Dis Sci.2020.

52. Jacobsen EA, et al. Lung Pathologies in a Chronic Inflammation Mouse Model Are Independent of Eosinophil Degranulation. Am J Respir Crit Care Med.2017;195:1321–1332.

53. Cardot-Ruffino V, et al. Generation of an Fsp1 (fibroblast-specific protein 1)-Flpo transgenic mouse strain. Genesis.2020;58:e23359.

54. Venkateshaiah SU, et al. Regulatory effects of IL-15 on allergen-induced airway obstruction. J Allergy Clin Immunol.2018;141:906–917 e906.

55. Zhang W, et al. S100a4 Is Secreted by Alternatively Activated Alveolar Macrophages and Promotes Activation of Lung Fibroblasts in Pulmonary Fibrosis. Front Immunol.2018;9:1216.

56. Harrop CA, Gore RB, Evans CM, Thornton DJ, Herrick SE. TGF-beta(2) decreases baseline and IL-13-stimulated mucin production by primary human bronchial epithelial cells. Exp Lung Res.2013;39:39–47.

57. Ingram JL, et al. Airway fibroblasts in asthma manifest an invasive phenotype. Am J Respir Crit Care Med.2011;183:1625–1632.

